# Diclofenac-eugenol modulation of Kv7 and TRPV1 ion channels

**DOI:** 10.1101/2024.03.16.585319

**Authors:** Adrián Rivera-Ruedas, Alejandro Rafael Medina-Vilchis, Juan José Romero-Tovar, Gema R. Cristóbal-Mondragón, Victor De la Rosa

## Abstract

Pharmacological targeting of ion channels represents a crucial avenue for pain management. Among these, the Kv7 family of ion channels plays a significant role controlling neuronal excitability and the generation and propagation of pain-related nerve impulses, thus mitigating excessive electrical signaling and curtailing the exaggerated transmission of pain signals. Pain management strategies often involve a multimodal approach, combining various medications with distinct mechanisms of action to achieve optimal outcomes. Eugenol possesses a spectrum of biological activities, including analgesic and anti-inflammatory properties. When used in conjunction with the anti-inflammatory drug diclofenac, eugenol demonstrates enhanced analgesic efficacy in animal models. We investigated the effects of diclofenac and eugenol on Kv7 and TRPV1 ion channels, both agents act as Kv7 activators, whereas diclofenac inhibits the TRPV1 current, and eugenol reduce the capsaicin-activated current presumably competing for the same binding site. Eugenol shows a time dependent biphasic effect on acid-activated TRPV1 current, first activating and then a slow decay of the current. When eugenol and diclofenac are used together, they limit the extent of depolarization of cells expressing Kv7 and TRPV1. Our results shed light on the combined effectiveness of eugenol and diclofenac in the treatment of acute pain.

## Introduction

A wide variety of molecules and proteins constitute the molecular mechanisms of pain. Several ion channels are functionally coupled either with receptors or enzymes that modulate them, or with other ion channels. Pharmacological regulation of these proteins is an obvious therapeutic target for pain treatment. The Kv7 family of ion channels form selective potassium pores in cell membranes. Their most relevant physiological contribution involves controlling membrane potential, the frequency of action potentials, and contributing to the phenomenon of adaptation, key factors in neuronal disorders and hyperalgesia [1]. These channels are present in sensory neurons and in pain transmission pathways in both the central and peripheral nervous systems [2, 3]. Their primary function in relation to pain is to regulate the excitability of neurons and control the generation and propagation of pain-related nerve impulses. This means they help prevent excessive generation of electrical signals and limit the exaggerated transmission of pain signals.

Multimodal treatment of pain involves using a combination of different therapies or approaches to manage pain effectively. This approach recognizes that pain is a complex experience influenced by various factors and addressing it through multiple methods can often lead to better outcomes compared to relying on a single intervention. Among the main strategies for pain relief is using a combination of medications with different mechanisms of action, this may include NSAIDs, opioids, anticonvulsants, antidepressants, or complementary and alternative medicines such as the use of molecules of natural origin.

Eugenol (4-hydroxy-2-methoxy phenol) is a naturally occurring phenolic compound found in essential oils of various plants, including cloves, nutmeg, and cinnamon. Eugenol possesses diverse biological properties such as analgesic, antioxidant, antiviral, and antibacterial properties [4-6]. It has also been studied by its anti-inflammatory, antidepressant, antiepileptic, neuroprotective, and anticancer activity [7-9]. Eugenol has been observed to affect both peripheral and central neurons, potentially affecting TRPV1, TRPV3 and TRPA1 channels [10-12]. Noteworthy, it has been reported that eugenol exhibits neuroprotective properties in the central nervous system after ischemia [13]. In the pharmaceutical realm, application include topical creams and emulsions for oral anaesthesia, it is often used as an ingredient in dental cements, temporary fillings, and impression materials, as it can have a sedative effect on the tissues in the oral cavity.

In animal models, the used of the anti-inflammatory drug diclofenac together with eugenol displays increased analgesic efficacy [14]. Although the main mechanism of action of diclofenac is related to COX blockade [15], there are also reports that it targets ion channels, including channels of the Kv7 family [16]. Our aim in this work was to determine the effect of diclofenac and eugenol on Kv7 and TRPV1 ion channels and analyse the effectiveness of both drugs in combination.

## Experimental procedures

HEK-293 cells were grown on tissue culture dishes in Dulbecco’s modified Eagle’s medium (Thermo Fisher Scientific) with 10% heat-inactivated fetal bovine serum (Gibco) plus 1% penicillin/streptomycin in a humidified incubator at 37 °C and 5% CO_2_ and passaged every 3-4 days using 0.05% of Trypsin-EDTA (Thermo Fisher Scientific) to lift the cells from the culture dish. cDNAs were transfected with Lipofectamine 2000 (Thermo Fisher Scientific) according to manufacturer’s instructions, experiments were performed 1 or 2 days after transfection. Cells were co-transfected with the indicated cDNA, together with EGFP cDNA as a transfection reporter.

Macroscopic currents were recorded using the whole cell or perforated patch clamp. Pipettes were pulled from borosilicate glass capillaries (BF150-86-10HP; Sutter Instruments) and fire polished with a microforge (Narishige). Membrane current was measured with an Axopatch 200A patch clamp amplifier and digitized for storage and analysis using a Digidata 1322A interface controlled by pClamp9.2. Series resistance was routinely compensated 60%, liquid junction potential corrections were not applied.

The external solution for Kv7 channels contained (in mM): NaCl 145, KCl 5, CaCl_2_ 2, MgCl_2_ 1, HEPES 10, pH7.4; HEPES was replaced with MES for acidic solutions and no CaCl_2_ was used for Kv7+TRPV1 co-transfection experiments. Internal solution contained (in mM): KCl 140, MgCl_2_ 2, EGTA 10, HEPES 10, GTP 2, ATP-Na_2_ 0.3, phosphocreatine 10, pH 7.3. TRPV1 bath and pipette solutions contained (in mM): 130 mM NaCl, 3 mM HEPES, 1 mM EDTA, pH 7.4. HEPES was replaced with MES for acidic solutions. A 1 M stock solution of Eugenol (Sigma) and 50 mM stock of Diclofenac (Sigma) were serially diluted in bath solution to the indicated concentrations, solution changes were accomplished by a gravity-fed superfusion to the recording chamber or by a fast-step perfusion system (Warner).

To estimate voltage dependence, tail current amplitudes (I_TAIL_) were measured after depolarizing voltages, normalized, and plotted as a function of test potential. I_TAIL_-V relations were fitted to a Boltzmann function of the form: 

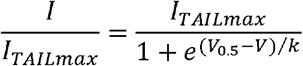

Where I_TAILmax_ is the maximum tail current, V_0.5_ is the voltage at which 50% of the conductance is reached and *k* is the slope factor. V_0.5_ values were plotted as a function of Eugenol concentration and fit to a Hill function of the form: 

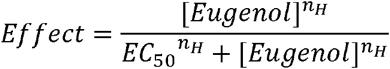

Where EC_50_ is the concentration at which 50% effect is reached and n_H_ is the Hill coefficient.

A 1 mM concentration of Eugenol was used in all other experiments. Data shown represents mean ± standard deviation for n determinations. Clampfit 10.6 and OriginPro 8.6 software were used for data analysis.

## Results and discussion

We tested the effect of eugenol on whole cell currents from HEK293 cells transiently expressing Kv7.2/3 channels (figure 1). Eugenol induces a mild increase of the outward current, and a more noticeable shift of the activation curve to hyperpolarization voltages, indicating that eugenol functions as an activator of the Kv7.2/3 current in a dose-dependent manner with an EC50 of 45 μM (figure 1D). Our recordings show an apparent increase in the rate of activation of the current, but further analysis of the activation time constant by fitting a single exponential to the second half of the rising phase of the current, shows that this is not the case (figure 1C). Therefore, eugenol appears to shift the activation voltage dependence without affecting other biophysical properties of the channels.

**Figure 1.**
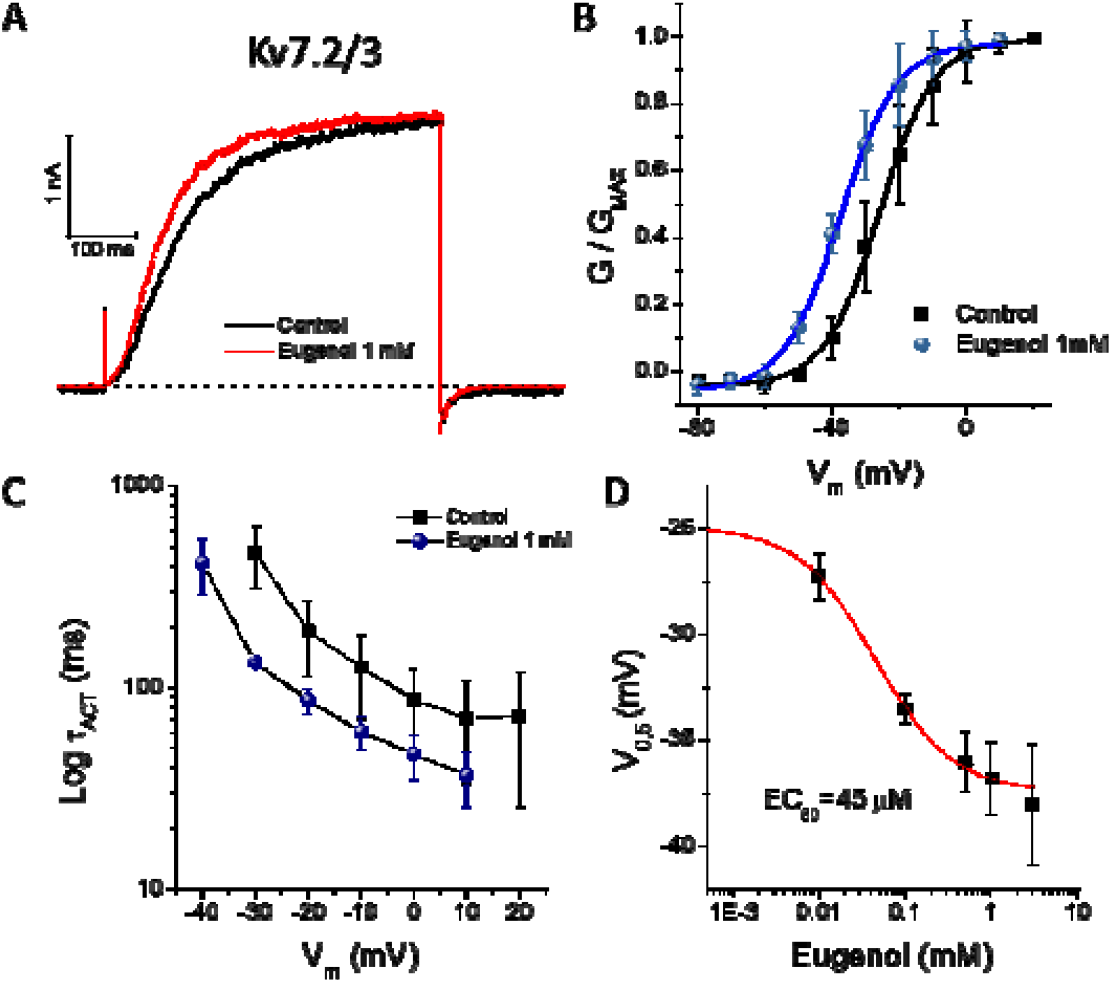
Eugenol activates the Kv7.2/3 ion channels. A) representative macroscopic M-currents recorded at a voltage step at 0 mV from a holding potential of -80 mV. B) normalized conductance in saline solution and in the presence of eugenol 1 mM, the maximum voltage shift was ∼15 mV. C) a single exponential was fitted to the second part of the current at the indicated voltage, eugenol did not change the activation kinetics. D) dose-response curve of the voltage shift by eugenol.

Next, we aimed to discriminate if eugenol affects both Kv7.2 and Kv7.3 similarly. We transfected each channel independently. Since Kv7.2 has less affinity to PIP_2_ [17], we noticed a markedly rundown of the current in the whole cell configuration, consequently we switch to perforated patch for these recordings that allowed us to minimize the rundown of the current to almost unnoticeable values during the time of the experiment. Eugenol activates the Kv7.2 channel and shifts the activation curve to hyperpolarizing voltages but to a lesser extent than in Kv7.2/3 (figure 2A and B). The shift in the activation was slightly noticeable and as shown in figure 2B, it could be masked by the standard error, therefore we ran ramp protocols to confirm the shift in the activation to more negative voltages. Correspondingly, eugenol activates the Kv7.3 channel in a dose dependent manner, the maximum voltage shift was 10 mV with an EC50 of 52 μM (figure 2C and D). For this set of experiments, we used the Kv7.3 (A315T) mutant, this channel shows larger whole cell current but the open probability and PIP_2_ sensitivity are the same as the WT channel [18, 19]. These results indicate that eugenol activates the so-called M current and shifts its voltage dependency to hyperpolarizing voltages. Most part of the shift in the activation can be explained by the effect on Kv7.3 channel, whereas both Kv7.2 and Kv7.3 contribute to the effect on the amplitude. Kv7 ion channel family is expressed in sensory neurons and is increasingly recognized as one of the important mechanisms to control nociceptive fiber activity [1]. The loss of function of these channels results in neuronal hyperexcitability. In nociceptive neurons, this is translated into increased excitability, nociception, and pain. Hence pharmacologic activation of the Kv7 channels could be analgesic.

**Figure 2.**
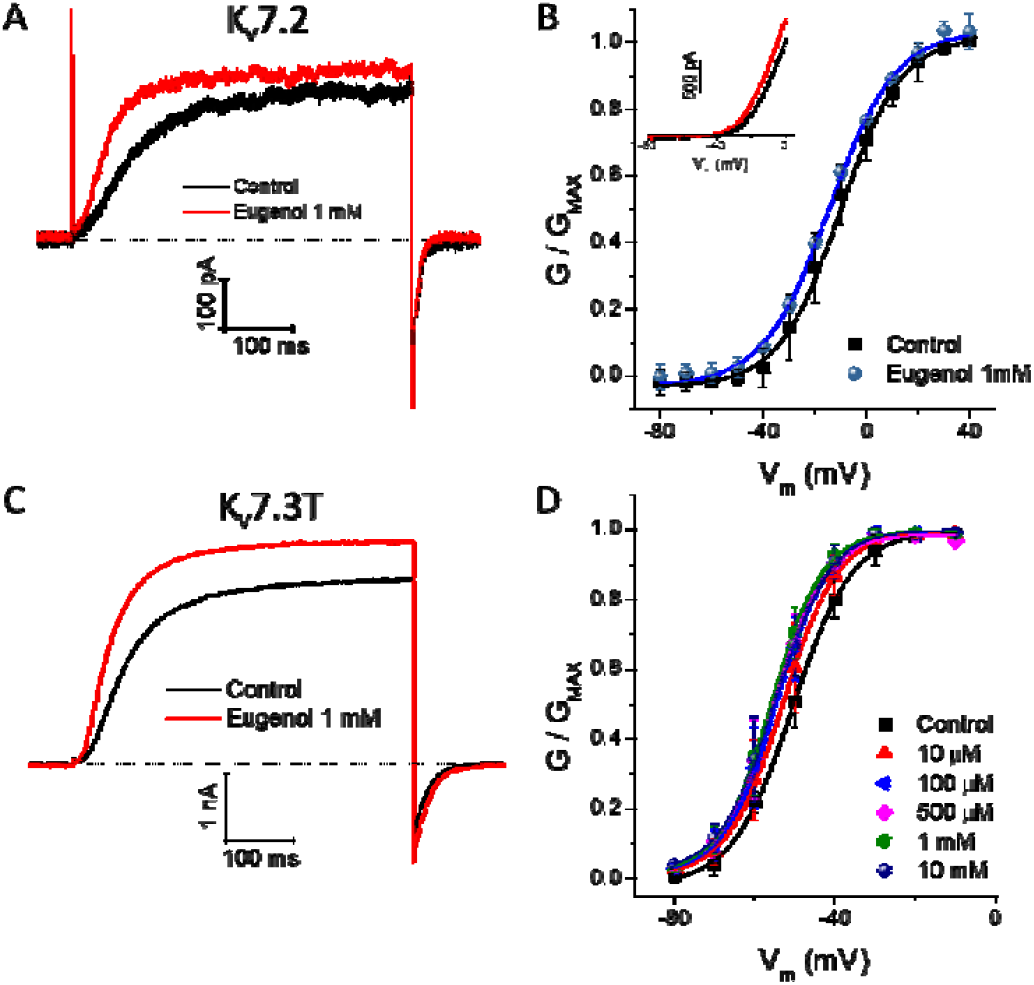
Differential contribution of the total activation of the M-current by eugenol. A and B) eugenol causes only a moderate hyperpolarizing shift on the voltage dependency of the Kv7.2 channels. The inset is the current in response to a voltage ramp from -80 mV to 0 mV. C and D) the voltage dependent shift by eugenol is principally caused by the effect on Kv7.3, recorded in the mutant Kv7.3 (A315T). The representative recordings were obtained by a voltage step to 0 mV from a holding potential of -80 mV.

Eugenol has been spotted as an activator of the TRPV1 channel [20], indeed it is a vanilloid derived from guaiacol, same as capsaicin, the widely studied TRPV1 activator. As reported earlier [12, 20], eugenol activates the TRPV1 channel but to a lesser extent than capsaicin (figure 3A and B), moreover when a non-saturating concentration of capsaicin is applied, eugenol decreases the whole cell current, indicating that both ligands share the same binding site. Recently, it was reported that the effects of eugenol on TRPV1 channels depends on the mode of activation of the channel [21], although we disagree with the interpretation of eugenol been an inhibitor of capsaicin-induced TRPV1 currents, we do recognize that when TRPV1 channels are activated with low pH, eugenol decreases the current (figure 3C). We differ with the interpretation of an open channel blocker suggested by Takahashi [21], our results indicate that after acidic activation of the channel, eugenol first increases the whole cell current, followed by a progressive reduction of the current with a time constant of 194 ±80 seconds. After reaching the maximum inhibition it was impossible to recover the current after washout of eugenol and keeping the cell at pH 7.4. Since our solutions have no calcium, we suggest that eugenol induces a calcium independent desensitization of TRPV1 when it is activated by acid, inhibiting their gating, and altering the physiological response to pain.

**Figure 3.**
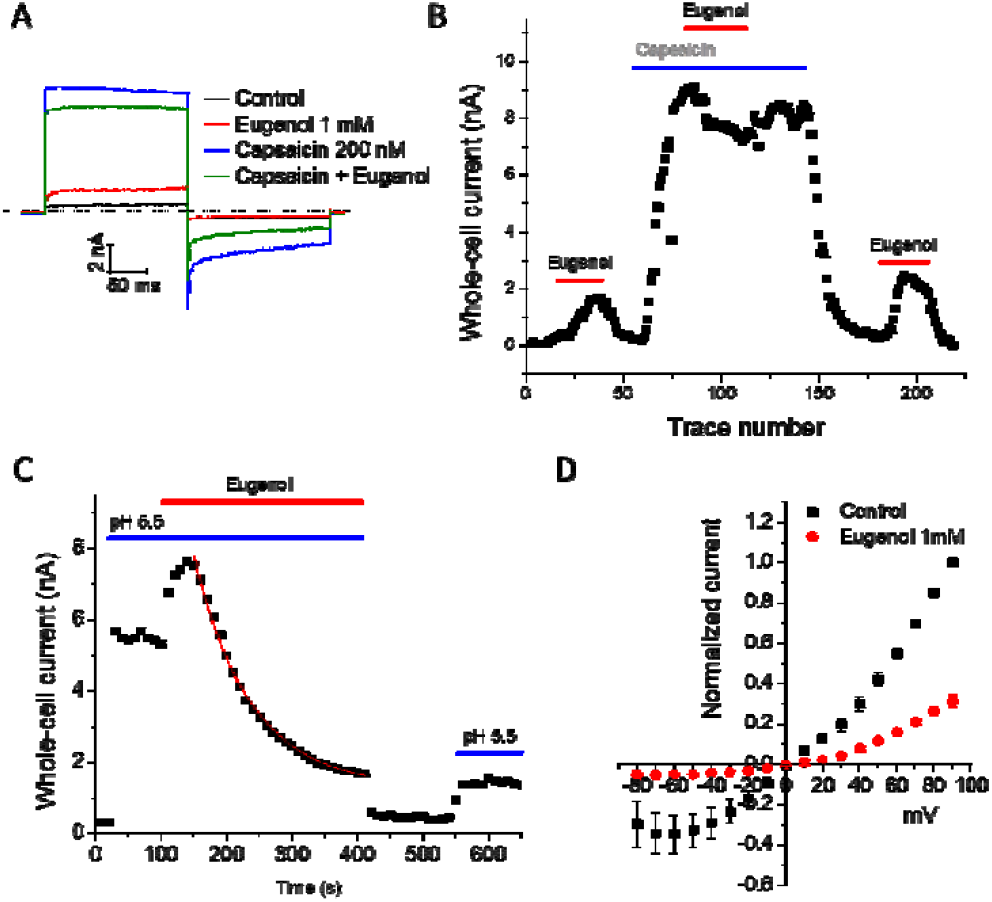
A) eugenol moderately activates the TRPV1 ion channel. In the presence of a non-saturating concentration of capsaicin, eugenol decreases the total current. Whole cell currents were recorded at a voltage step of 80 mV followed by a step to -80 mV from a holding potential of 0 mV. B) temporal course of the experiment shown in A. C) the acid-dependent TRPV1 current is activated and then inhibited in the presence of eugenol. The red line is an exponential fit of the decay of the current, giving on average a 194 second decay. D) comparison of the current vs voltage curves in saline and after inhibition of the current by eugenol.

The nonsteroidal anti-inflammatory drug, diclofenac also serves as an activator of the Kv7.2/3, 7.4 and a blocker of Kv7.5 channels [16, 22]. In our hands, we did not observe a significant increment or a hyperpolarizing shift at 50 μM of diclofenac in Kv7.2/3 as reported earlier [16], instead the activation was only observed at 100 μM (figure 4A and B). Also at the same concentration, diclofenac inhibits the TRPV1 currents (figure 4C). No further concentrations were tested.

**Figure 4.**
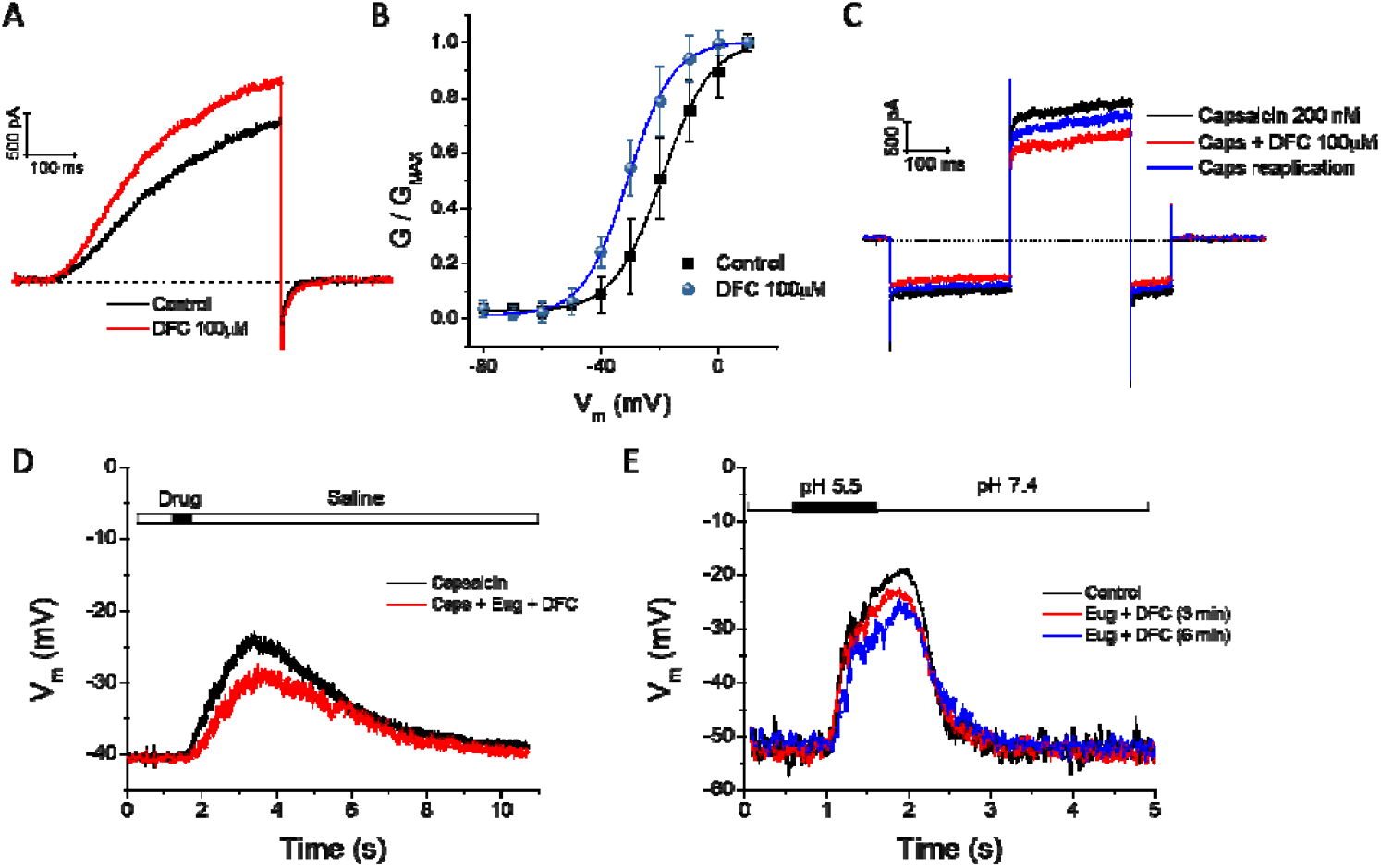
A) representative recordings of the M-current before and after application of diclofenac. B) voltage shift to hyperpolarizing potentials of the activation of the M-current caused by diclofenac. C) at non saturating concentrations of capsaicin, diclofenac inhibits the TRPV1 current. D) current-clamp recordings of HEK293 cells co-transfected with TRPV1 and Kv7.2/3 channels. Application of capsaicin at the time indicated by the black line in the top, induces a depolarization of the membrane. The depolarization is less pronounced when capsaicin is applied in combination with eugenol and diclofenac. E) current-clamp recording similar to panel D, where the depolarization was induced by application of an acidic solution. The cells were held at -60 mV for 3 or 6 minutes, where cells were constantly perfused by eugenol + diclofenac. After the time intervals, the acidic solution was again applied. The depolarization was less after perfusion of the drugs.

The activation of Kv7.2/3 together with the inhibition of TRPV1 by eugenol and diclofenac may contribute to explain the synergistic interaction between these drugs in the treatment of acute pain [14]. To test this possibility, we co-transfected TRPV1 and Kv7.2/3 channels and measured the depolarization of the cells with capsaicin or acidic pH and in the presence of eugenol + diclofenac. As shown in the figure 4D, cell membrane depolarization caused by capsaicin is less when both eugenol + diclofenac are present, similarly, cell membrane depolarization caused by acidic pH is less after continuous perfusion of eugenol + diclofenac (figure 4E), these results indicate that these drugs can limit the extent of depolarization and consequently the number of action potentials, reinforcing the analgesic potential of the drugs when applied in combination. Although we did not try the drugs directly on sensory neurons, it is known that capsaicin sensitive dorsal root ganglion neurons express Kv7.2, Kv7.3 and Kv7.5, in fact, retigabine, a Kv7 agonist, reportedly reduced noxious heat responses, while XE991, a Kv7 antagonist, increased the noxious responses [23]. Thus, the reports indicate that M channels strongly affect the excitability of nociceptive fibers [1, 23, 24], and our results give additional supporting evidence on this subject.

It has been previously shown that administration of diclofenac induces an increase in the expression of IL-1β between 3 and 12 hours after administration[25], in the case of eugenol + diclofenac there was also an increase in IL-1β but the dose of the drugs was 20 times lower [26]. Our results shown here are focused on the acute antinociceptive efficacy of both drugs in combination. Furthermore, during inflammation, an acidic microenvironment modulates proinflammatory and anti-inflammatory responses, hence, it is possible that there is an acid dependent activation of TRPV1 that ultimately induces thermal hyperalgesia. As shown in our results, the acid dependent activation of TRPV1 is greatly inhibited by eugenol, this effect together with the activation of Kv7.2/3 channels, can contribute to explain the acute analgesia observed by these drugs.

## ACKNOWLEDGEMENTS

The work was supported by grant FORDECYT-PRONACES/1308052/2020 from CONACyT, Mexico.

## References

1. Du, X., et al., M-type K(+) channels in peripheral nociceptive pathways. Br J Pharmacol, 2018. 175(12): p. 2158–2172.

2. Passmore, G.M., et al., KCNQ/M currents in sensory neurons: significance for pain therapy. J Neurosci, 2003. 23(18): p. 7227–36.

3. Wladyka, C.L. and D.L. Kunze, KCNQ/M-currents contribute to the resting membrane potential in rat visceral sensory neurons. J Physiol, 2006. 575(Pt 1): p. 175–89.

4. Guenette, S.A., et al., Pharmacokinetics and anesthetic activity of eugenol in male Sprague-Dawley rats. J Vet Pharmacol Ther, 2006. 29(4): p. 265–70.

5. Jorkjend, L. and L.A. Skoglund, Effect of non-eugenol- and eugenol-containing periodontal dressings on the incidence and severity of pain after periodontal soft tissue surgery. J Clin Periodontol, 1990. 17(6): p. 341–4.

6. Zhang, P., et al., Enhanced chemical and biological activities of a newly biosynthesized eugenol glycoconjugate, eugenol α-d-glucopyranoside. Applied Microbiology and Biotechnology, 2013. 97(3): p. 1043–1050.

7. Anjum, N.F., et al., Novel derivatives of eugenol as potent anti-inflammatory agents via PPARγ agonism: rational design, synthesis, analysis, PPARγ protein binding assay and computational studies. RSC Adv, 2022. 12(26): p. 16966–16978.

8. Fangjun, L. and Y. Zhijia, Tumor suppressive roles of eugenol in human lung cancer cells. Thorac Cancer, 2018. 9(1): p. 25–29.

9. Fujisawa, S. and Y. Murakami, Eugenol and Its Role in Chronic Diseases, in Drug Discovery from Mother Nature, S.C. Gupta, S. Prasad, and B.B. Aggarwal, Editors. 2016, Springer International Publishing: Cham. p. 45–66.

10. Chung, G., et al., Activation of transient receptor potential ankyrin 1 by eugenol. Neuroscience, 2014. 261: p. 153–60.

11. Bandell, M., et al., Noxious cold ion channel TRPA1 is activated by pungent compounds and bradykinin. Neuron, 2004. 41(6): p. 849–57.

12. Xu, H., et al., Oregano, thyme and clove-derived flavors and skin sensitizers activate specific TRP channels. Nat Neurosci, 2006. 9(5): p. 628–35.

13. Won, M.H., et al., Postischemic hypothermia induced by eugenol protects hippocampal neurons from global ischemia in gerbils. Neurosci Lett, 1998. 254(2): p. 101–4.

14. González-Lugo, O.E., et al., Synergistic interaction between 4-allyl-1-hydroxy-2-methoxybenzene (eugenol) and diclofenac: An isobolograpic analysis in Wistar rats. Drug Dev Res, 2020.

15. FitzGerald, G.A. and C. Patrono, The coxibs, selective inhibitors of cyclooxygenase-2. N Engl J Med, 2001. 345(6): p. 433–42.

16. Peretz, A., et al., Meclofenamic acid and diclofenac, novel templates of KCNQ2/Q3 potassium channel openers, depress cortical neuron activity and exhibit anticonvulsant properties. Mol Pharmacol, 2005. 67(4): p. 1053–66.

17. Hernandez, C.C., B. Falkenburger, and M.S. Shapiro, Affinity for phosphatidylinositol 4,5-bisphosphate determines muscarinic agonist sensitivity of Kv7 K+ channels. J Gen Physiol, 2009. 134(5): p. 437–48.

18. Choveau, F.S., et al., Phosphatidylinositol 4,5-bisphosphate (PIP2) regulates KCNQ3 K(+) channels by interacting with four cytoplasmic channel domains. J Biol Chem, 2018. 293(50): p. 19411–19428.

19. Choveau, F.S., et al., Pore determinants of KCNQ3 K+ current expression. Biophys J, 2012. 102(11): p. 2489–98.

20. Yang, B.H., et al., Activation of vanilloid receptor 1 (VR1) by eugenol. J Dent Res, 2003. 82(10): p. 781–5.

21. Takahashi, K., T. Yoshida, and M. Wakamori, Mode-selective inhibitory effects of eugenol on the mouse TRPV1 channel. Biochem Biophys Res Commun, 2021. 556: p. 156–162.

22. Brueggemann, L.I., et al., Diclofenac distinguishes among homomeric and heteromeric potassium channels composed of KCNQ4 and KCNQ5 subunits. Mol Pharmacol, 2011. 79(1): p. 10–23.

23. Passmore, G.M., et al., Functional significance of M-type potassium channels in nociceptive cutaneous sensory endings. Front Mol Neurosci, 2012. 5: p. 63.

24. Du, X., et al., Control of somatic membrane potential in nociceptive neurons and its implications for peripheral nociceptive transmission. Pain, 2014. 155(11): p. 2306–22.

25. Barcelos, R.P., et al., Diclofenac pretreatment modulates exercise-induced inflammation in skeletal muscle of rats through the TLR4/NF-κB pathway. Appl Physiol Nutr Metab, 2017. 42(7): p. 757–764.

26. González-Lugo, O.E., et al., ANALGESIC ED50 DICLOFENAC-EUGENOL COMBINATION EFFECT ON PROINFLAMMATORY CYTOKINES – IL-1β, IL-6, TNFα AND MAPKs – IN RAT MUSCLE. Farmacia, 2022. 70(4): p. 583–588.

